# Functionally Convergent B Cell Receptor Sequences in Transgenic Rats Expressing a Human B Cell Repertoire in Response to Tetanus Toxoid and Measles Antigens

**DOI:** 10.1101/159368

**Authors:** Jean-Philippe Büerckert, Axel R.S.X. Dubois, William J. Faison, Sophie Farinelle, Emilie Charpentier, Regina Sinner, Anke Wienecke-Baldacchino, Claude P. Muller

## Abstract

The identification and tracking of antigen-specific immunoglobulin (Ig) sequences within total Ig repertoires is central to high-throughput sequencing (HTS) studies of infections or vaccinations. In this context, public Ig sequences shared by different individuals exposed to the same antigen could be valuable markers for tracing back infections, measuring vaccine immunogenicity, and perhaps ultimately allow the reconstruction of the immunological history of an individual. Here, we immunized groups of transgenic rats expressing human Ig against tetanus toxoid (TT), Modified Vaccinia virus Ankara (MVA), measles virus hemagglutinin and fusion proteins expressed on MVA and the environmental carcinogen Benzo[a]Pyrene, coupled to TT. We showed that these antigens impose a selective pressure causing the Ig Heavy chain (IgH) repertoires of the rats to converge towards the expression of antibodies with highly similar IgH CDR3 amino acid sequences. We present a computational approach, similar to differential gene expression analysis, that selects for clusters of CDR3s with 80% similarity, significantly overrepresented within the different groups of immunized rats. These IgH clusters represent antigen-induced IgH signatures exhibiting stereotypic amino acid patterns including previously described TT and measles specific IgH sequences. Our data suggest, that with the presented methodology, transgenic Ig rats can be utilized as a model to identify antigen-induced, human IgH signatures to a variety of different antigens.

## Introduction

Immunoglobulin (Ig) molecules are the primary effectors of the humoral immune response. In theory, Ig can bind to every possible antigen through the large variety of immunoglobulin V (variable), D (diversity) and J (joining) gene rearrangements in the bone marrow and target-oriented affinity maturation in germinal centers (1). All B cells of a germinal center are clonally related to a common ancestor and target the same antigen with varying affinities, iteratively selecting for improved affinity and avidity (2). The Ig molecules of the emerging B cells bind to the target epitope in a lock-and-key principle which is mediated mainly by the heavy chain complementary-determining region 3 (CDR3) loop on top of the Ig (1, 3). The CDR3 is the most variable part of the Ig sequence and the main antigen-binding determinant. The repertoire of CDR3s sufficiently describes the entire functional immunoglobulin heavy chain (IgH) repertoire of an individual (3, 4).

High-throughput sequencing (HTS) has been widely applied to study the IgH repertoire in response to vaccination and infection (5). With this technique, it has become possible to investigate the evolutionary affinity maturation processes after antigenic challenge and to compare their outcome across individuals (6). The IgH repertoire is essentially private (7), but it appears that individuals also produce a public response to a common antigenic stimulus characterized by a certain degree of similarity at the CDR3 sequence level (8–11). Studies investigating this concept of public Ig CDR3s mainly used human blood-derived PBMCs. These represent only a miniscule part of the complete Ig repertoire (12) and it is critical to capture the affinity-matured B cells during their brief transit from the germinal centers through peripheral blood to the bone marrow. Public CDR3s were identified in humans in response to dengue infection, H1N1 seasonal influenza vaccination and repetitive polysaccharide antigens (5, 11, 13). Such CDR3s provided signatures of past immunological exposures allowing for sequence-based monitoring of vaccination or infectious diseases, and perhaps ultimately to reconstruct an individual’s antigenic history.

Here we applied HTS on bone marrow B cells rich in serum antibody producing plasma cells from rats carrying human germline IgH and light chain (IgL) loci, the OmniRat™ (14–17). These transgenic rats were immunized with viral (Modified Vaccinia virus Ankara, MVA), protein (measles virus hemagglutinin and fusion proteins, HF and Tetanus toxoid, TT) and chemically defined hapten-conjugate antigens (Benzo[a]Pyrene-TT, BaP-TT) to study the evolution of convergent CDR3 amino acid sequences. We showed that OmniRat™ mount convergent Ig responses characterized by CDR3s with high amino acid sequence similarity. The level of similarity was consistent for all investigated antigens. We applied an approach similar to differential expression analysis to identify overrepresented clusters of highly similar, antigen-driven CDR3s. These could be grouped into antigen-associated signatures matching previously described measles virus hemagglutinin-specific OmniRat™ hybridomas (18) and human TT-specific antibodies (13, 19–21). Our results suggested that humanized Ig transgenic rats can be used as a model to study human-like Ig repertoire dynamics and to determine antigen-associated CDR3 signatures to characterize the history of antigen exposure in human individuals.

## Materials and Methods

### Animals and immunizations

Humanized Ig transgenic rats (OmniRat™, Open Monoclonal Technology Inc., Palo Alto, USA) were developed and bred as previously described (14–17). Animals were separated into 6 groups of 4 to 6 individuals. They received 3 intraperitoneal injections at 2-weeks intervals and were sacrificed 7 days after the last injection. Injections either contained 100 μg of tetanus toxoid (TT group; Serum Institute of India, Pune, IN) or of a benzo[a]pyrene-TT conjugate construct (BaP-TT group) (22), both formulated with 330 μg of aluminum hydroxide. Other rats were injected with 10^7^ PFU of a recombinant Modified Vaccinia virus Ankara (MVA) expressing the hemagglutinin (H) and fusion (F) glycoproteins of the measles virus (MVA-HF group) or the MVA viral vector only (MVA group) without adjuvant. The control animals received either 330 μg of aluminum hydroxide alone (ALUM group) or were left untouched (NEG group). Antigen-specific IgG responses were monitored by ELISA 10 days after immunizations and at sacrifice. All animal procedures were in compliance with the rules described in the Guide for the Care and Use of Laboratory Animals (23) and accepted by the ‘Comité National d’Éthique de Recherche’ (CNER, Luxembourg).

### Antigens for immunization and ELISA

BaP was coupled to ovalbumin (OVA, Sigma-Aldrich) for ELISA and to purified tetanus toxoid as previously described (22). The recombinant Modified Vaccinia virus Ankara (MVA) and the recombinant MVA carrying measles virus H and F proteins of the Edmonston strain (MV vaccine strain, clade A) viruses were propagated on BHK-21 cells (ATTC™ CCL-10™) as previously described (24–26). Antigen-specific IgG antibody levels in sera were determined in 384-well microtiter plates (Greiner bio-one, Wemmel, BE), coated overnight at 4°C with either 250 ng of MV antigen (Measles grade 2 antigens, Microbix Biosystems, Mississauga, USA), 2.5 × 10^5^ PFU of sonicated MVA (∼314 ng), 187.5 ng of TT or 0.25 μM of BaP-OVA in carbonate buffer (100 mM, pH9.6). Free binding sites were saturated with 1% bovine serum albumin (BSA) in Tris-buffered saline at room temperature for 2h. Serial dilutions of the sera were added for 90 min at 37°C, and developed with alkaline phosphatase-conjugated goat anti-rat IgG (1/750 dilution, ImTec Diagnostics, Antwerp, BE) and the appropriate substrate. Absorbance was measured at 405 nm. Endpoint titers (EPT) were determined as the serum dilutions corresponding to 5 times the background.

### Sample preparation, amplification and IonTorrent PGM Sequencing

Lymphocytes were isolated from bone marrow samples by density-gradient centrifugation (ficoll^®^ Paque Plus, Sigma-Aldrich). Total RNA was extracted from 10^8^ cells with an RNeasy midi kit following the manufacturer’s protocol (Qiagen) and enriched for mRNA using paramagnetic separation (μMACS mRNA Isolation kit, Miltenyi Biotech, Leiden, NL). cDNA was prepared from 300ng of mRNA using dT18 primers and Superscript III reverse transcriptase (Thermo Fisher Scientific) at 50°C for 80 min. Recombined IgH fragments were subsequently amplified by multiplex PCR using primers for human IgHV region and rat Cγ region with Q5 Hot Start High Fidelity polymerase (NEB, Ipswich, USA) as described previously (18). Amplicons were size selected on a 2% agarose gel and quantified. Quality was checked with a Bioanalyzer (High Sensitivity DNA, Agilent Technologies, Diegem, BE). Four randomly-selected libraries were pooled in equimolar concentrations and sequenced on a 318™ Chip v2 (Thermo Fisher Scientific) using multiple identifiers (MIDs) with the Ion OneTouch™ Template OT2 400 Kit and the Ion PGM Sequencing 400 Kit (Thermo Fisher Scientific) on the Ion Torrent Ion Personal Genome Machine (PGM™) System (Thermo Fischer Scientific).

### Quality control and sequence annotation

BAM files were extracted from the Torrent Suite ™ software (version 4.0.2, standard settings) and demultiplexed by multiplex identifiers (MID). Only reads with an unambiguously assigned MID (0 mismatches), identified primers at both ends (2 mismatches allowed) and more than 85% of the bases with a quality score above 25 were considered for further analysis. After clipping MIDs and primers, sequences were collapsed and submitted to the ImMunoGeneTics database (IMGT) HighV-QUEST webserver (www.imgt.org, (27)) for IgHV gene annotation and CDR3 delineation (28). IgHV and IgHJ genes for the in-frame, productive sequences were subsequently assigned using a local installation of IgBlast (29), including only the genes present in the genome of the OmniRat™ as references. Only sequences with an unambiguously assigned IgHV and IgHJ gene were considered for further analysis.

### CDR3 similarity threshold for public immune responses

The number of matches for the 200 most frequent CDR3s (top 200) of a rat A in a rat B was obtained for a series of similarity thresholds and returned as ratio from 0 to 1 (i.e. all top 200 CDR3s of rat A have a match in rat B). Ratios were determined from 50% to 100% sequence similarity in one percent increments. The averages for the top 200 matching ratios at each increment were then calculated for all rats within a vaccination group and all rats vaccinated with unrelated antigens. Rats with related antigens were excluded in the pairwise comparison (e.g. MVA as intra-group for MVA-HF). The average top 200 matching ratios were plotted against sequence similarity along with the first derivatives in GraphPad Prism 5 (www.graphpad.com).

### Identification of antigen-driven CDR3 clusters

Only CDR3s longer than 4 amino acids were included in the analysis. CDR3s with a minimum of 80% amino acid similarity, were considered as relatives. One amino acid length difference was allowed, to account for insertion and deletions introduced by SHM and occasional differential CDR3-IgHJ region alignments by IMGT (30, 31). These length difference was penalized the same way as a substitution. For each CDR3, the cumulative count of all its 80% relatives per rat (CDR3-count) was calculated and stored in a fuzzy match count table. Data was imported and analyzed with DESeq2 according to the standard workflow for RNA-seq, treating CDR3-counts as expression values (32). Briefly, data was imported as a count-data matrix and converted into a DESeq2-object with conditions according to the antigens used for vaccination. Correct sample grouping was confirmed using variance stabilizing transformed count data (VST-counts). Euclidian distance computation was performed on VST-counts as described in the DESeq2 vignette (33). Principle component analysis plots were generated using the ‘PlotPCA’ function on VST-counts of the DESeq2 package. P-values were adjusted for multiple testing and to determine the false discovery rate (FDR) using Benjamini-Hochberg correction (34). Based on an FDR of 1%, over-represented CDR3 sequences were extracted if their adjusted p-values were lower than 0.01. Log2-fold change cut-offs were determined manually per antigen group. The extracted CDR3 sequences were grouped using single-seed iterative clustering based on maximum difference of 80% sequence similarity. All analytical scripts were written in Python 2.7 and R 3.2.3 (35).

### 3D modeling

Selected Ig nucleotide sequences were uploaded to IMGT for annotation. Sequences were elongated to full length by adding the missing nucleotides from the closest germline gene as predicted by the IMGT algorithm. Full length sequences were submitted to the “Rosetta Online Server that Includes Everyone” (ROSIE, http://rosie.rosettacommons.org/, (36–38)) with enabled H3 loop modeling option. ROSIE-output PDB files of the grafted and relaxed models were visualized using PyMol (version 1.7.4, http://pymol.org, (39)).

## Results

### High-throughput sequencing of OmniRat™ IgH mRNA transcripts

To study convergent IgH repertoires in response to vaccination, 32 transgenic IgHumanized rats (OmniRat™) were immunized with different antigens (Table 1; TT, BaP-TT, MVA, MVA-HF). Aluminum hydroxide (ALUM) was used as an adjuvant for TT and BaP-TT. Two control groups received either the adjuvant alone or were left untouched (NEG). All animals exhibited a specific antibody response against the immunizations and mock immunized (ALUM group) and non-immunized animals (NEG group) showed no detectable antigen-specific antibodies (Supplemental Fig. S1). MVA-HF and BaP-TT vaccinated animals exhibited a specific immune response against the MVA vector or the TT carrier protein respectively, albeit at lower levels than the animals immunized with these antigens only (Supplemental Fig. S1). Rearranged heavy chain IgG genes were amplified from mRNA extracted from bone marrow (BM) lymphocytes and sequenced on a high-throughput sequencing (HTS) Ion Torrent PGM™ platform. A total of 37,473,982 raw reads with MID were obtained (range: 850,298 – 1,879,372 per animal, Supplemental Table S1). After quality control and annotation, on average 86,619 unique nt sequences per animal were retained for analysis. The rats expressed a diverse IgH repertoire, including varying frequencies of all human IgHV and IgHJ genes. All possible IgHVJ combinations were found in all vaccination groups with no bias in IgHV, IgHJ genes or IgHVJ recombination usage.

**Table 1.**
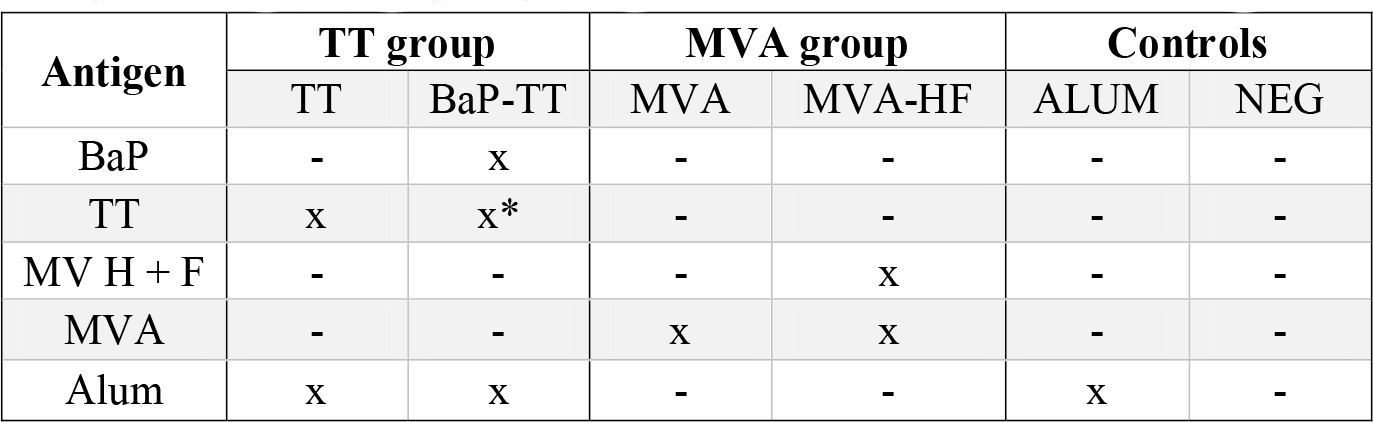
Study design: antigen and vaccination groups. TT used in the BaP-TT vaccination group was chemically modified during the coupling process and therefore does not present the same antigenic surface as native TT in the TT group.

### Unique and highly similar CDR3 sequences in response to the same antigen

We first investigated to what extent rats that received the same antigen expressed the same CDR3 amino acid sequences. Pairs of rats from different vaccination groups (369 pairs) shared less CDR3s with each other than pairs of rats within the same group (71 pairs, p-value < 2×10^−16^, Kruskal-Wallis with Nemenyi post-hoc test) or immunized with related antigens (56 pairs, p-value = 3.4×10^−14^), indicating that mutual CDR3s are essentially induced by the immunizations (Fig. 1). Among a total of II, 643 identical CDR3s (i.e. 100% similarity) that were shared by any set of two or more rats irrespective of the antigen, 5,346 CDR3s (45.9%) were shared exclusively by animals of the same group and 1,912 (16.4%) were shared between animals immunized with a related antigen (TT and BaP-TT, MVA and MVA-HF). Most of the CDR3s shared within groups were common to only two animals of the same group (6,467; 89.1% of CDR3s shared within groups only). CDR3s present in all animals of a group were rare (Table 2). However, multiple CDR3s which differed only by one or two amino acids were shared by all animals within a vaccination group but not by animals from other groups (Table 3). Interestingly, these differences occurred preferentially at certain positions of a CDR3 amino acid sequence. This suggested that the vaccinations seemed to have induced identical CDR3s as well as clusters of highly similar CDR3s.

**Figure 1.**
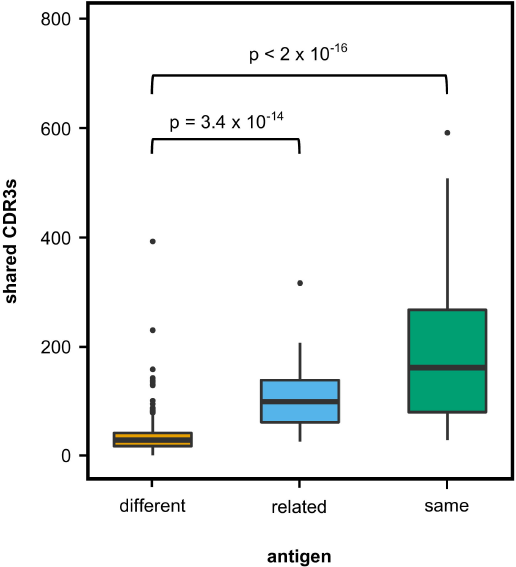
Shared CDR3s in OmniRat™ pairs. Box-whisker plots represent the number of identical CDR3s shared between pairs of rats from different (369 pairs, orange), related (56 pairs, light blue), or the same vaccination group (71 pairs, green). More CDR3s were shared between rats from the same (p-value 3.4 × 10^−14^) or related (p-value < 2 × 10^−16^) antigen group than between rats of different antigen groups (Kruskal-Wallis test followed by Nemenyi post hoc test).

**Table 2:**
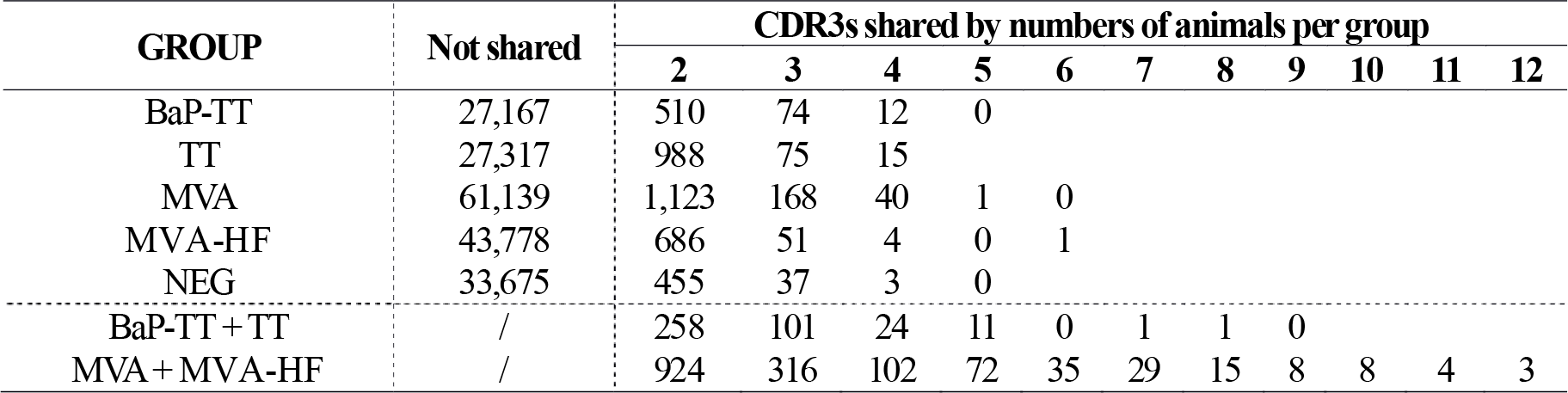
CDR3s shared between rats in the same vaccination group

**Table 3:**
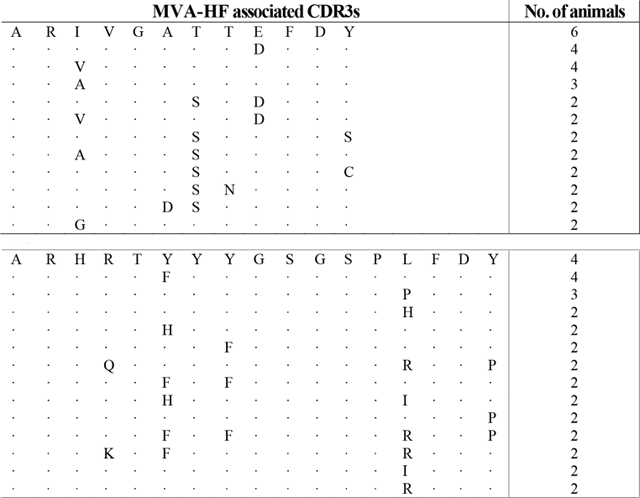
Selected CDR3 sequences shared by rats in the MVA-HF group only

### Shared antigen-related CDR3s at 80% sequence similarity

We compared CDR3s within and across the different vaccination groups to estimate the degree of similarity between these antigen-related clusters. We determined which of the top 200 CDR3s of any rat A had a related CDR3 in a rat B either within the same group (intra-group comparison) or between groups (inter-group comparison) allowing for a single amino acid substitution. The same analysis was repeated for two, three and up to eight amino acid substitutions. The number of top 200 CDR3s found to be present inter- and intra-group were plotted against the amino acid substitutions expressed as percent of CDR3 length (Fig. 2A, Supplemental Fig. S2). The resulting sigmoidal curves showed a similar shape for all vaccination groups. In the exponential phase between 100% and 90-95%, intra-group overlap was higher than inter-group overlap. In the linear phase between 90-95% and 75%, overlap increased faster for the inter-group comparison. In the asymptotic phase below 75% similarity, both inter- and intra-group overlap leveled off towards 1, indicating that all top 200 CDR3s of a rat had relatives in any other rat, irrespective of the antigen administered. The first derivatives of the curves showed that in all cases the inflection point was at around 80% (Fig. 2B, Supplemental Fig. S2). Thus, at this similarity threshold a maximum number of related CDR3s can be found within the same group while keeping the number of related CDR3s between groups at a minimum. In conclusion, all antigens induced in these rats a public IgH response that can best be characterized by clusters of CDR3s with at least 80% similarity.

**Figure 2.**
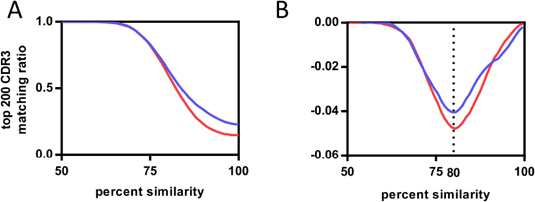
Influence of CDR3 sequence similarity on CDR3 repertoire overlap between rats. (A) Average fractions of top 200 CDR3s of the MVA-HF vaccination group shared with all CDR3s of other samples. Samples were divided into two groups having either the same antigen (HF group samples, blue curve) or different antigens (ALUM, BaP-TT, TT and NEG samples, red curve). Both curves follow a similar sigmoidal behavior. (B) First derivative of both curves. Inflection points align at 80% CDR3 amino acid similarity.

### Hierarchical clustering of CDR3 repertoires at 80% sequence similarity

Based on the above observation, we identified antigen-driven CDR3s using a workflow developed for differential gene expression analysis of RNA-seq data (32). For each CDR3 within a rat, counts of CDR3 sequences with 80% similarity (CDR3-counts) were used analogous to RNA-seq read counts. Rats of the same vaccination group were considered as replicates. The CDR3-counts followed a negative binomial distribution (Fig. 3A). Compared to RNA-seq data, CDR3s usually lack a baseline expression and are essentially private, resulting mostly in zero-counts for individuals across the study, while some shared CDR3s have very high counts in a single animal (Fig. 3B). To account for this distribution, we applied variance stabilizing transformation (VST) to the CDR3-counts reducing the variance of the standard deviations over ranked means (Fig. 3C). Hierarchical clustering of VST-counts revealed three clusters (Fig. 3D). Cluster I included all animals immunized with MVA (with or without MV HF protein expression). Interestingly, within this cluster, animals of the MVA-HF group and of the MVA group emerged from two separate branches indicating, that additional presentation of HF antigens leaves a distinct imprint in the CDR3 repertoire. Cluster II contained the three groups of animals that received alum as an adjuvant (TT, BaP-TT and ALUM). Again, each of the three groups clustered on separated sub-branches. Cluster III contained only untreated animals (NEG group) and was distinct from all immunized animals. The low variance and the specific grouping of the samples through both principle components showed that the VST-counts cluster the data by vaccination group. This indicated that the different antigens had distinct impact on a subset of the Ig repertoire of the rats (Fig. 3E). When the data were reanalyzed applying an 85% or 75% threshold, the clear clustering of rats by vaccination group was lost (**Supplemental Fig. S3**), thus confirming that the 80% similarity threshold was optimal to identify antigen-associated responses on the CDR3 repertoire of the rats. Additionally, it showed that VST CDR3-count data can be analyzed analogously to RNA-seq count data.

**Figure 3.**
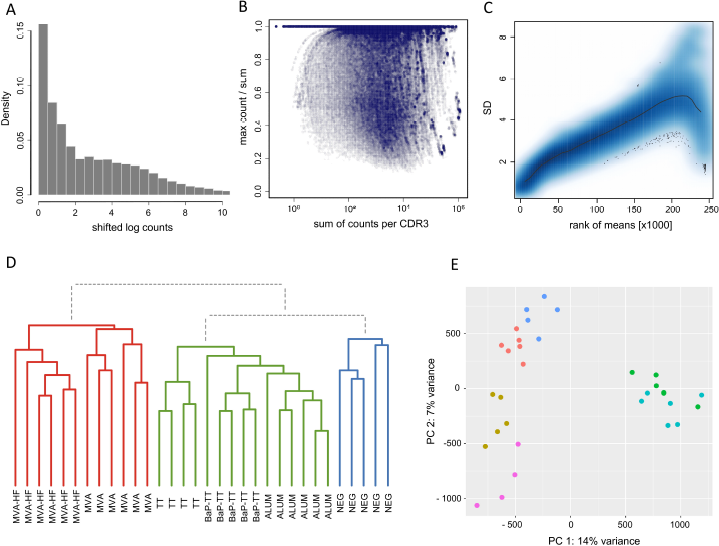
DESeq2 statistics and sample grouping. (A) Density-histogram representing the distribution of CD R3-counts (log(x+1) transformed). The CD R3-counts follow a negative binomial distribution (B) Sparsity-plot displaying the count distribution per CDR3. The sum of counts for every CDR3 is plotted in log_10_-scale against the highest count for the CDR3 divided by the sum of all counts for the CDR3. Density of data is indicated by hue. (C) CDR3-wise standard deviation of ranked means of counts after variance stabilizing transformation (VST-counts). The black line shows the standard deviation for all ranked means of VST-counts across all samples, the blue area indicates the data distribution and density by hue. (D) Dendrogram of the Euclidian sample distances calculated for VST-counts. Three main clusters are indicated by coloration (Cluster I: red, Cluster II: green, Cluster III: blue). (E) Scatterplot for the first two principal components of VST-counts. Samples are colored by vaccination-group (MVA: light blue, MVA-HF: green, TT: pink, BaP-TT: gold, ALUM: red, NEG: blue).

### Large numbers of antigen-associated CDR3s group into stereotypic signatures

Similar to RNA-seq expression experiments, we aimed to identify CDR3s that are differentially represented between groups of rats. Based on a false discovery rate (FDR) of 1%, 16,727 of the 249,657 (6.9%) unique CDR3s across all groups were found to be overrepresented. One hundred-fold differences in numbers of overrepresented CDR3s were identified in each of the six antigen-groups (Table 4). The highest number of overrepresented CDR3s was found in the two combined groups MVA and MVA-HF (n=11,080, 10.4% of the unique CDR3s for this combined group), and TT and BaP-TT (n=2,451, 4.4%), which reflected the high immunogenicity of the antigens TT and MVA common within these groups. Less overrepresented CDR3s were found in the MVA (n=1,689, 2.6%), the MVA-HF (n=804, 1.8%) and TT group (n=540, 1.8%). The lowest number of overrepresented CDR3s was found in the BaP-TT group (n=163, 0.6%). These overrepresented CDR3s could be considered group-specific, and thus immunization induced.

**Table 4:**
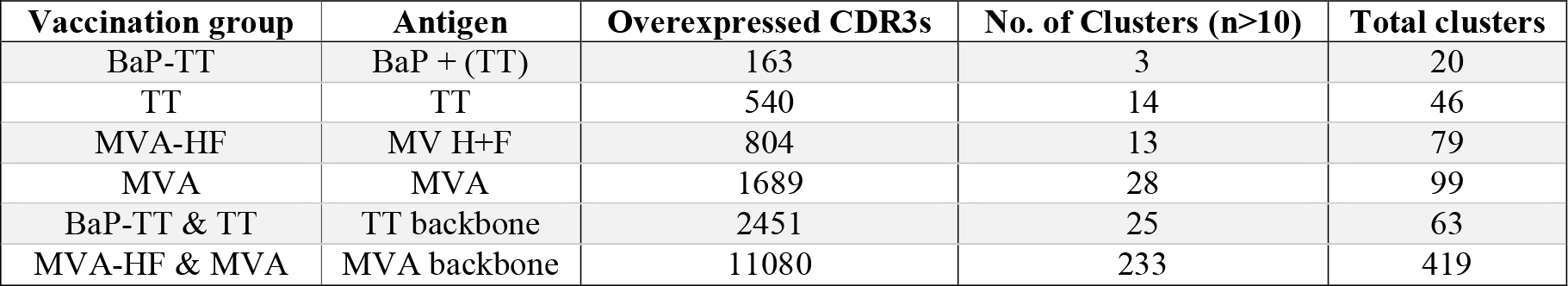
Antigen-driven sequences and 80% similarity clusters

Overrepresented CDR3s were grouped into clusters of 80% sequence similarity (Fig. 4). The larger the antigen, the more clusters were found. For instance, 20 clusters were found for the BaP-hapten while 109 clusters were found for the TT protein (46 for TT alone, and 63 for TT and BaP-TT combined). The largest number of clusters was found for the MVA virus antigen (518, with 99 for MVA alone and 419 for MVA and MVA-HF combined). These complex antigen-driven clusters of CDR3s, typical for each group, represented up to 46.5% of the bone marrow IgH repertoire of the rats (Fig. 5). The fraction of the repertoire corresponding to these CDR3 clusters varied between the groups but was relatively consistent among animals of the same antigen-group. All together we showed that OmniRat™ exhibited large fractions of highly similar, stereotypic CDR3s in response to the applied vaccinations, even across groups with shared antigens.

**Figure 4.**
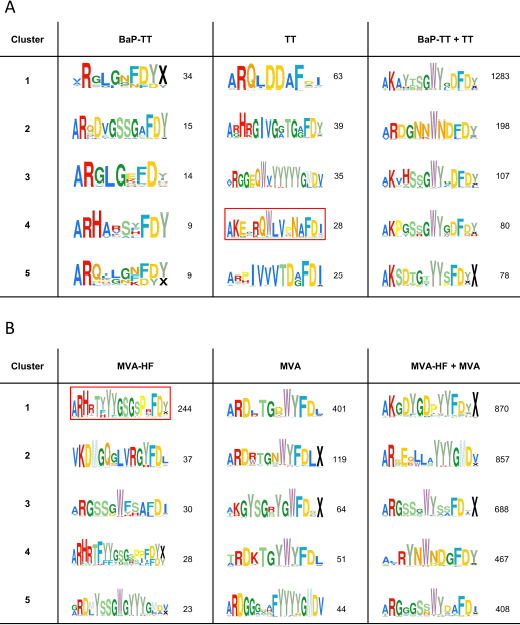
Antigen-associated CD R3-similarity clusters. The top 5 clusters of 80% similar CDR3s overexpressed in response to the antigens are shown as Weblogos. Coloration follows IMGT amino acid coloration scheme (54). Numbers represent the unique CDR3s in each cluster. (A) Clusters associated with the antigens BaP-TT, TT and the combined antigen groups BaP-TT and TT. The red box indicates TT-associated OmniRat™ CDR3s bearing an amino acid pattern also found in human anti-TT PBMC CDR3 sequences from for independent studies. (B) Clusters associated with the antigens MVA-HF, MVA and the combined antigens MVA and MVA-HF. The red box indicates MV-HF associated CDR3-signature also identified in OmniRat™ hybridomas generated in an independent experiment in response to MV antigens.

**Figure 5.**
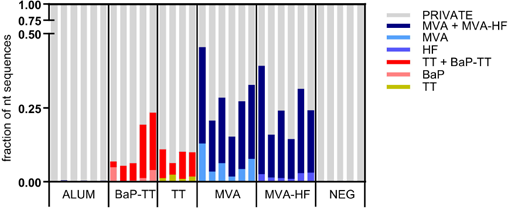
Fractions of the nucleotide Ig repertoire encoding for CDR3 signatures. The Ig repertoire per sample is displayed using numbers of full length nucleotide sequences. Nucleotide sequences encoding for CDR3s that are part of a signature are colored by associated antigen.

### Stereotypic signatures match known MV-specific and TT-specific CDR3s

MVA-HF signatures were compared to the previously described CDR3s of MV-specific hybridoma clones derived from an independent set of OmniRat™ immunized with whole MV antigen (18). The largest of the identified HF associated clusters (244 members) matched three CDR3s of MV-specific hybridoma cells, suggesting that this cluster is an MV-H or F protein induced CDR3 signature (Fig. 6). Similarly, our TT-associated clusters were compared to known human TT-specific IgH sequences (13, 19–21). The CDR3s from the TT-associated *cluster 4* matched 12 published human CDR3s (Fig. 7A). This OmniRat™ CDR3 signature as well as the human CDR3s consisted of 15-mer CDR3s following the same amino acid pattern. Both humans and rats elicited a conserved paratope defined by a static motif ‘+QWLV’ (+= R/K) at the center of the CDR3, flanked by variable positions that are connected to the torso of the CDR3 (Fig. 7B). This indicates that similar key positions are used even across species. The sequence similarity between the human and rat CDR3s ranged from 67% to 87% resulting from different torso amino acid compositions at the positions flanking the conserved binding motif (Fig. 7C+D). To compare the structures of these CDR3s from human and rat origin, we performed 3D-homology modeling on their Fab-fragments. Four human antibodies with available heavy and light chain sequences (20) and four selected OmniRat™ heavy chain sequences paired with the human light chains were modeled with Rosetta Antibody. Within the OmniRat™-human chimeric Fab-fragments, the CDR3s formed torso structures ranging from unconstrained amino acid formations over short beta-sheets to rigid beta-sheet hairpin constructs (Fig. 7C). Like the rats, human CDR3s exposed the key binding residues at the very tip of the CDR3 loop by a rigid beta-sheet hairpin formation of the torso that protruded out of the IgH core structure (Fig. 7D). Together our results corroborate the evolution of functionally convergent CDR3s in different individuals and by different vaccines delivering the same antigen. Also, this strongly indicates that OmniRat™ and humans, albeit the lower sequence similarity between their TT-associated CDR3s, produce antibodies with highly homologous CDR3s in response to the same antigen.

**Figure 6.**
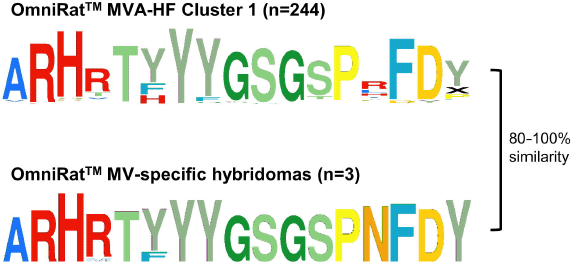
OmniRat™ MV-specific CDR3 signature. The clusters of CDR3s overrepresented in response to MVA-HF (see also Fig. 4A) and the CDR3s from 3 monoclonal hybridomas specific for MV proteins are shown as Weblogos. The differences between the sequences were calculated as Levenshtein distances in percent of CDR3 length.

**Figure 7.**
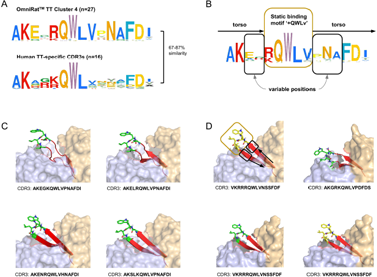
Omnirat™ and human antibodies against tetanus toxoid with similar properties and structures. (A) Sequence similarity range between OmniRat™ TT-associated *cluster 4* (Fig. 4B) CDR3s and human TT-specific CDR3s (Levenshtein distance as percent of sequence length). (B) Amino acid pattern for the combined TT-specific human and TT-associated rat CDR3s. The Weblogo shows the conserved binding motif ‘+QWLV’ (+ indicates K/R), and torso amino acids with variable positions are highlighted. (C) 3D-homology models of four OmniRat™-HC-human-LC chimeric antibody Fab-fragments. Heavy chains are colored in orange and light chains in blue, both visualized with 50% transparent surface. CDR3 torsos are shown as cartoon and colored in red. Binding motifs are displayed as sticks and colored in green with only polar hydrogens shown. Views were enlarged to focus on the CDR3 structure. (D) 3D-homology models of four human Fab-fragment visualized as described for (C). Motif, variable and torso structures are highlighted with boxes and arrows as described in (B).

## Discussion

We analyzed more than 2,700,000 functional IgH sequences derived from the bone marrow of transgenic rats expressing human B cell receptor genes immunized with different antigens. Our study showed that these rats produced identical as well as highly similar CDR3 amino acid sequences in response to common antigenic challenges. When shared CDR3 repertoire fractions were investigated at different levels of sequence similarity, overlaps between rats from the same vaccination group were optimal around 80% CDR3 amino acid similarity. Applying a differential gene expression workflow to the counts of 80% similar CDR3s, we presented a novel way to identify convergent, stereotypic CDR3 sequences in response to an antigenic stimulus. These included known CDR3s induced by different measles antigens, indicating that the identified CDR3s are specific for the measles virus H or F proteins which were shared in both immunizations. In addition, our approach also identified CDR3s in response to tetanus toxoid that were remarkably similar to known tetanus-specific CDR3s from human samples. Our findings highlighted the presence of convergent IgH transcripts at high levels in the bone marrow of the transgenic rats and that these sequences are highly similar to those of humans.

Pairs of rats within the same group shared more identical CDR3s than pairs from different groups, but very few CDR3s were shared among all rats of a group. Given the tremendous size and diversity of the Ig repertoire, finding identical sequences in several individuals is indeed unlikely (40). Because of private processes during B cell development including stochastic affinity maturation of the Ig molecules, a certain variability in CDR3s converging towards reactivity with the same antigen is to be expected (6, 41, 42). Galson *et al.* found that for the identification of public repertoires in humans an 87.5-91.6% cut-off (1 in 12 to 1 in 8 amino acids) was optimal to identify TT- and influenza-related CDR3 clusters (43). In the present study, we explored the relation between CDR3 sequence similarity and the overlap between CDR3 repertoires, by inter- and intra-group cross-comparisons at different levels of sequence similarity. Our data showed that 80% amino acid similarity optimized the intra-group overlap between CDR3 repertoires while keeping the inter-group overlap at a minimum. The identified antigen-associated clusters were absent in rats outside immunization groups, which provides a strong support for their underlying biological relevance.

We showed that between 6% and 46% of the bone marrow IgH repertoire correspond to convergent CDR3 sequences. Similar proportions (15-50%) of antigen-specific CDR3 sequences were reported in peripheral blood B cells of patients with acute dengue infections (5). In contrast, in the reconvalescent dengue patients as well as in influenza patients, convergent sequences represented only to less than 1% of peripheral B cell sequences. Such human studies are normally restricted to peripheral blood where only a small fraction of the repertoire can be found and assessed (12). In mice, elevated levels (37%) of public clones were observed in response to HBsAg and less to NP-HEL (22%) and OVA (14%) when examining bone marrow derived long-lived plasma cells (CD138^+^ CD22^−^ MHCII^−^ CD19^−^ IgM^−^PI^−^) (44). Here, we analyzed Ig mRNA from bulk rat bone marrow cell isolates, an organ rich in serum antibody-producing long-lived plasma cells (45). Because long-lived plasma cells with high levels of Ig mRNA and only class switched BCR (IgG) were targeted, those are overrepresented in our datasets (42, 43). This explains the high levels of converging CDR3s in the bone marrow. We thus primarily target antigen-associated effector B cells, facilitating the tracking of antigen-specific sequences induced by similar antigens.

Potential influence on the repertoire composition could also result from PCR amplification biases introduced during library preparation as well as sequencing errors (46). We did not account for potential errors and sequencing bias by using molecular barcodes or similar methods (47–50). However, the data analysis of the present study was based on collapsed, unique nucleotide Ig sequences, minimizing the influence of potential PCR amplification bias. Analysis of the nucleotide sequences before and after collapsing to unique nucleotide sequences revealed no major difference in our findings, indicating that PCR amplification bias did not falsifying our results. The IonTorrent PGM sequencing platform is prone to insertion and deletion errors, especially within homopolymer repeats (51). Such errors cause frameshifts within the Ig sequence which are detected by IMGT with 98% efficiency in a benchmarking setup, missing only indels at the beginning and end of the sequence or if placed in close proximity to each other masking the resulting frameshift (Bürckert *et al*., manuscript in preparation). Sequences with detected indels are marked by IMGT as productive with detected errors and were not included in the described analysis. Furthermore, our analysis is based on the CDR3 amino acid sequence. An insertion or deletion within the CDR3 encoding nucleotides results in the sequence being labelled as unproductive, with no correction attempts undertaken by IMGT (Bürckert *et al*., manuscript in preparation). Less than 1% of indel combinations remain undetected by IMGT and could be present within the CDR3 encoding nucleotides (Bürckert *et al*., manuscript in preparation). These rare combination of sequencing errors would then result in artefactual CDR3s either covered by the applied 80% sequence similarity clustering threshold or missed because of higher sequence variation. Therefore, such CDR3 artefacts can be expected to induce only a small underrepresentation of CDR3s by lowering CDR3-counts. In conclusion, the presented workflow is well protected from potential sequencing errors or PCR bias, that could impact our conclusions.

We found that certain CDR3s have high counts of 80% relatives within a group but very few to none in the unrelated groups. This is in principle comparable to differential gene expression in RNA-seq data. The CDR3-counts followed a negative binomial distribution but, unlike in RNA-seq experiments, our data contained large amounts of CDR3s with zero-counts over different samples. These correspond to private CDR3s that are absent in other rats of the same or other groups. On the other hand, some CDR3s exhibited very high counts of 80% relatives within an animal. While such a data distribution is uncommon in RNA-seq, they were nevertheless compatible with our computational approach (DESeq2, (32)) as demonstrated by negative binomial data distribution and perfect sample grouping after variance stabilizing transformation of the CDR3-counts. Interestingly, the Euclidian distance grouping of MVA-HF and MVA rats remained unchanged for 85% and even for 75% CDR3-counts in contrast to TT-associated rats. Similarly, Trück *et al.* found highly similar (< 2 mismatches) Hib- and TT-related sequences enriched seven days post-vaccination, but could not identify H1N1- and MenC-related sequences at the same threshold (13). Correspondingly, statistical evidence of convergent CDR3s in pairs of donors against influenza with a mean genetic distance of ∼75% were reported (52). Together with our data this indicated that the identification of convergent Ig repertoire responses using amino acid similarity thresholds was applicable. Future research will tell to what extent the 80% threshold can be applied to other antigens.

Identified convergent CDR3 matched to sequences of previously described human monoclonal antibodies against TT protein (13, 19-21). Despite the relatively low sequence similarity (67% to 87%) between OmniRat™ and human TT-specific CDR3s, they shared a common sequence and structural motif at the center of the CDR3. The center part of the CDR3 is exposed at the tip of the loop structure which directly interacts with the antigen while the adjacent amino acids act as a supporting scaffold. Similarly, Victor Greiff and coworkers observed stereotypical motifs at the center of the CDR3 amino acid sequences in specific antibodies following NP vaccination in mice (53). While structural similarity cannot readily be used to determine antibody specificity, algorithms to identify convergent CDR3s could be further improved by including structural parameters drawn from the expanding amount of available crystal structures.

In conclusion, we demonstrated a strong public IgH response with converging and overlapping CDR3 repertoires in animals exposed to the same antigens. These converging repertoires consisted of similar CDR3 sequences that can be best described using an 80% amino acid similarity threshold. Additionally, we presented an approach to identify such CDR3s by adopting a group-wise expression analysis, similar to RNA-seq approaches. This provides also a valuable tool for large-scale HTS data-mining to identify potential candidates for high-affinity targeted antibody design.

## Author’s contributions

JB and AD contributed equally to the work. JB designed and developed the bioinformatics approach, interpreted data, performed data processing and wrote the manuscript. AD designed and carried out research, prepared samples, interpreted data and wrote the manuscript. WF supported bioinformatics approaches and data processing, corrected the manuscript. AB set up the raw data processing script and performed data processing, data interpretation and corrected the manuscript. SF, EC provided technical assistance with immunizations, ELISA and virus culture. RS performed IonTorrent PGM sequencing. CM designed research, interpreted data, corrected the manuscript and supervised work. All authors have read and approved the final version of the manuscript.

## Acknowledgements

We are grateful to R. Buelow from Open Monoclonal Technology Inc. (Palo Alto, CA, USA) for providing the OmniRat™ We thank Dr. B. Moss, NIAID, National Institutes of Health (Bethesda, USA) for providing the MVA and recombinant MVA viruses. We thank Josiane Kirpach for her valuable discussions and Fleur AD Leenen for critically revising the manuscript.

## Abbreviations

Abbreviations used in this article:

ALUM: aluminum hydroxide
BaP: Benzo[a]Pyrene
BaP-TT: Benzo[a]Pyrene tetanus toxoid conjugate
BM: bone marrow
EPT: endpoint titers
F: fusion glycoprotein of the measles virus
FDR: false discovery rate
H: hemagglutinin glycoprotein of the measles virus
HF: hemagglutinin and fusion protein of the measles virus
HTS: high-throughput sequencing
IgL: immunoglobulin light chain
IMGT: ImMunoGeneTics database
MV: measles virus
MVA: Modified Vaccinia virus Ankara
ROSIE: rosetta online server that includes everyone
TT: tetanus toxoid
VST: variance stabilizing transformation

